# The human language processing system straightens natural speech

**DOI:** 10.64898/2026.06.30.735613

**Authors:** Jiaming Xu, Tien Dung Nguyen, Jerry Tang, Alexander G. Huth, Robbe L. T. Goris

## Abstract

Large language models trained on next-word prediction have impressive linguistic capabilities. This suggests that the goal of temporal prediction is essential to language processing, but how this goal impacts the structure of speech representations in the human brain remains unknown. Here, we test the hypothesis that prediction is facilitated by the temporal straightening of representational trajectories along the speech processing hierarchy. We developed a methodology for measuring the curvature of these trajectories using fMRI. Our method exploits a previously unknown connection between the timescale of single-unit responses and the curvature of population trajectories. We examined brain responses of subjects listening to natural speech. Response trajectories were most curved in lower-level auditory areas and progressively straightened along the cortical hierarchy. We presented the same speech stimuli and perturbed versions thereof to wavLM—a speech representation model that is well aligned with human brain responses—and found that hierarchical straightening effects are strongest for stimuli whose statistical structure resembles natural speech. Together, our results establish a direct connection between the goal of temporal prediction, the geometry of neural speech representations, and the cortical hierarchy of representational timescales.

---

Prediction has emerged as a central, but controversial, framework for understanding human speech processing [1, 2, 3, 4, 5, 6, 7]. Speech is intrinsically structured in time and is therefore partially predictable. The idea that the brain constantly attempts to predict future events before they occur is attractive because it offers a unifying, efficient account of brain function. Yet, how the goal of temporal prediction might shape computation and representation in the human language processing system is an open question. Speech comprehension depends on a hierarchy of computations distributed across the cortex (Fig. 1a), from acoustic analysis in early auditory regions to phonological, lexical, and semantic processing in temporal, parietal, and frontal areas [8, 9, 10, 11, 12, 13, 14]. Early predictive coding theories posit that higher-level areas provide individual lower-level neurons with predictions about upcoming feedforward input. These predictions are compared with the actual input, leaving the neurons to explicitly represent prediction error [15, 16, 17, 18]. However, evidence for explicit predictive coding in language is limited [4]. Recent predictive coding theories instead propose that neural representations reflect the goal of prediction in a more implicit manner, by using prediction as a learning objective [19, 20, 21]. An example of such an implicit strategy in the domain of language is offered by Large Language Models (LLMs) trained on next-word prediction [22, 23]. In these models, each word is gradually transformed from its input form into a prediction of the next word. The sequence of transformations implicitly supports prediction by creating internal representations that facilitate temporal prediction [24, 22, 25, 26, 27].

**Figure 1.**
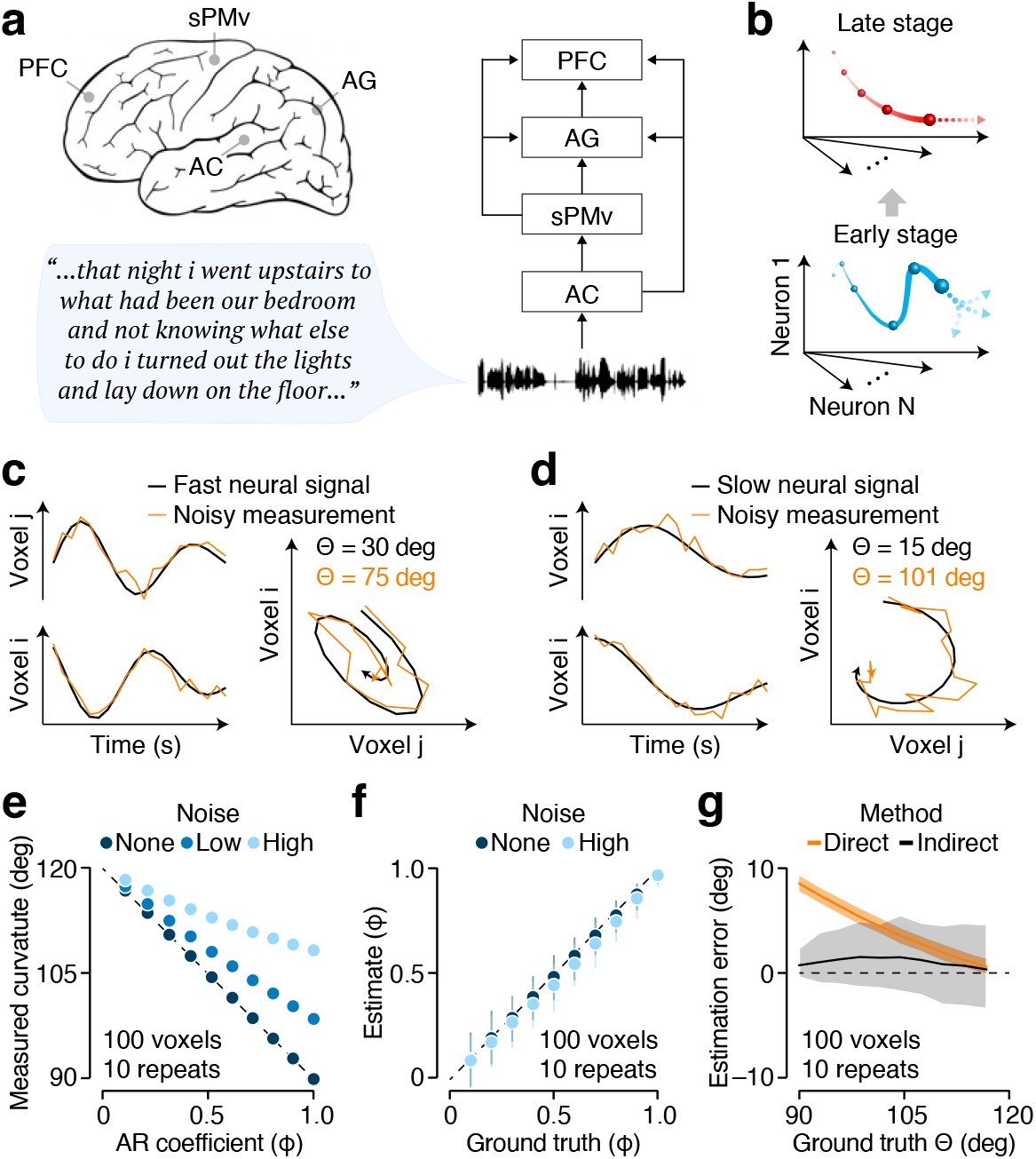
Estimating neural curvature from fMRI data: experimental design, computational hypothesis, and analysis methods. (**a**) Experimental design. fMRI BOLD responses were recorded while subjects listened to narrative stories (data originally reported in Ref. [30]). Our analysis is largely focused on four hierarchically organized areas in the human speech processing system. AC: Auditory cortex, sPMv: superior ventral premotor cortex, AG: angular gyrus, PFC: prefrontal cortex. (**b**) Computational hypothesis. The temporal straightening hypothesis posits that neural trajectories become straighter along the processing hierarchy in order to facilitate prediction of future internal states. (**c**) Response trajectories for a pair of units with fast-changing responses (black curve: neural signal; orange curve: noisy measurement). Noise added to the trajectory biases its curvature upward. (**d**) The response trajectory of a pair of units with slow-changing responses. Trajectory curvature is similarly biased upwards when noise is added. (**e**) Direct curvature estimates are plotted against the AR coefficient (which controls representational timescale) for a set of simulated 100-D trajectories, which varied in their signal (X-axis) and noise (color) properties. At low noise, the AR coefficient and curvature are tightly linked. Symbols show the average across 500 runs of the simulation. (**f**) Estimated representational timescale plotted against ground truth value for AR(1) response trajectories that varied in timescale (abscissa) and noise level (light vs dark colors). Symbols show the mean across 500 runs, error bars indicate the cental 95 % of the distribution.(**g**) Curvature estimation error as a function of ground truth value for two estimation methods (orange vs black), applied to a set of simulated 100-dimensional trajectories. Estimates are based on trial-averaged responses across 10 repeated presentations. Thick lines show the mean across 500 runs of the simulation, colored bands indicate the central 95 % of the distribution.

From a geometrical perspective, speech prediction is challenging because the timecourse of the raw sensory input is jagged (i.e. very different from timepoint to timepoint) and difficult to extrapolate. A natural solution is to transform this input into internal representations that follow straighter temporal trajectories, thus enabling the prediction of future internal states through linear extrapolation (Fig. 1b). This implicit strategy, known as ‘temporal straightening’, was first proposed as a computational mechanism employed by the primate visual system [19, 28]. Recently, it was demonstrated that temporal straightening also occurs in LLMs trained on next-word prediction. As language information propagates along the network hierarchy, representational trajectories gradually become straighter from the first to the middle layers of the network [25]. In a different strand of research, these same LLMs have been found to be much more effective at explaining brain activity elicited by natural speech than earlier models [29]. Together, these findings motivate the hypothesis that the human language processing system may exploit temporal straightening to support speech prediction. Here, we directly test this hypothesis for the first time in an fMRI experiment in which subjects listened to narrative stories (data originally reported in Ref. [30]).

To test the temporal straightening hypothesis for language in fMRI data, we first established a theoretical relationship between the geometry of population trajectories and a well-studied property of neural responses, the representational timescale [31, 32, 33]. For the simplest recurrent random process, i.e. a first-order autoregressive (AR) model, trajectory curvature is fully determined by the representational timescale. Hierarchical straightening thus implies a hierarchical organization of representational timescales. This insight naturally integrates our hypothesis about representational geometry with existing knowledge about representational timescales [31]. It also provides the basis for an indirect measure of curvature from fMRI data, which overcomes the noise-driven biases inherent in direct curvature estimation. As hypothesized, we found that response trajectories gradually straighten along the language processing hierarchy. To test the specificity of these effects, we additionally examined trajectory curvature in WavLM, a leading model for human cortical speech representations [34]. We found that hierarchical straightening effects are stronger for natural human speech than for perturbed control stimuli. Interestingly, the relationship between representational timescale and trajectory curvature was also evident in WavLM and in simulated populations of spiking neurons. Together, these findings establish temporal straightening as a representational motif in the human speech processing system, and demonstrate that there is a deep connection between the goal of temporal prediction, the geometrical structure of neural speech representation, and the hierarchical organization of representational timescales in language processing systems.

## Results

### Estimating neural curvature from fMRI data

We analyzed fMRI data collected while subjects listened to continuous narrative stories and compared the curvature of response trajectories across several brain regions involved in language processing. At each moment in time, the auditory input elicits a specific pattern of population activity within a given brain region. This pattern can be summarized as a single point in a high-dimensional space, whereby each unit in the population constitutes one dimension. As the narrative unfolds, these points trace out a trajectory that reflects the evolving population response. A natural measure for the local curvature of this trajectory is the unsigned angle between consecutive segments [19]. The average of these angles over time provides a direct measure of the curvature of an entire story-trajectory in each brain region. Intuitively, when population responses change rapidly, the resulting trajectory undergoes abrupt turns and is therefore highly curved (Fig. 1c, black curves). In contrast, when responses evolve more slowly, the trajectory changes direction gradually and is thus straighter (Fig. 1d, black curves). This suggests a connection between neural curvature [28] and representational timescale [31, 32, 33]: a population with short representational timescale should have higher curvature, while one with long representational timescale should have lower curvature. Such a connection would provide a framework for integrating hypotheses about representational geometry with established knowledge about representational timescales.

To test the idea that curvature and timescale are related, we considered response time series arising from a first-order autoregressive (AR(1)) model, the simplest recurrent random process. In an AR(1) model, each point in time is linearly dependent on the previous time point plus a random term: *y*_*t*_ = *ϕy*_*t*−1_ + *ϵ*_*t*_, where *y*_*t*_ is the response at time *t, ϵ*_*t*_ is a Gaussian random variable, and *ϕ* ∈ [−1, 1] is the autoregressive coefficient. This coefficient controls the timescale of dependencies in the resulting time series. We found an analytic relationship between populations of autoregressive responses (i.e. *n* AR(1) processes with the same *ϕ*, giving **y**_*t*_ ∈ ℝ^*n*^) and *θ*, the average curvature across the trajectory, which is given by *θ* = arccos((*ϕ* − 1)*/*2) (see Methods for derivation). This theoretical insight connects single unit properties to population geometry, but also implies that trajectory curvature can be inferred from the temporal dynamics of responses. Exploiting this relationship could thus help us overcome the well-known challenge of measuring curvature in the presence of noise [19, 28] (Fig. 1c,d, orange vs black curves).

We investigated the effect of noise on population curvature estimates by simulating AR(1) processes with noise comparable to that found in fMRI data (Methods). We varied *ϕ* to create trajectories that differed in ground-truth curvature (Fig. 1e, dark symbols). In the presence of noise, direct curvature estimates systematically deviated from the ground truth, with larger noise levels producing greater bias (Fig. 1e, light symbols). One way to mitigate measurement noise is to collect multiple trials using the same stimulus and average the responses. In our simulation, averaging across 10 trials reduced the bias, but did not eliminate it (Extended Data Fig. 1a). These simulations demonstrate that direct curvature estimates are strongly affected by measurement noise. The exact spatiotemporal structure of fMRI noise is difficult to estimate and likely differs across brain regions [35]. Therefore, interpreting cross-region differences in direct curvature estimates is fraught with difficulty.

To overcome this issue we developed a noise-robust procedure for estimating curvature indirectly via timescale. We first estimate a brain region’s timescale by fitting an AR(1) model to the voxel responses using ordinary least squares, and then use the analytic relationship between *ϕ* and *θ* to obtain an indirect estimate of response trajectory curvature (Methods). This procedure is robust to noise because AR(1) coefficients can be recovered reliably in the presence of fMRI-like measurement noise (Fig. 1f; Extended Data Fig. 1b). We compared direct and indirect curvature estimates in simulated data. Even at the single-trial level, indirect estimates systematically outperformed direct estimates, albeit with a small bias (Extended Data Fig. 1c). Trial-averaged responses (10 repeats) yielded nearly unbiased curvature estimates for the indirect method, but not for the direct method (Fig. 1g). Together, these results indicate that indirect curvature estimates based on timescale enable meaningful comparisons across brain regions.

### Indirect curvature estimates decrease along the processing hierarchy

We used the direct and indirect procedures to estimate curvature from fMRI data collected while subjects listened to natural speech [30]. We first focused on four regions of interest (ROIs) that span the language processing hierarchy: auditory cortex (AC), which processes acoustic and phonemic information; the superior ventral premotor area (sPMv; aka area 55b [36]), a mid-level language area in premotor cortex; angular gyrus (AG), a high-level language area [9]; and prefrontal cortex (PFC), another area processing high-level language information [9]. Consider the results for an example subject. Direct curvature estimates were uniformly high (all above 100°) and exhibited no obvious cross-area structure (Fig. 2a, orange symbols). This pattern is reminiscent of the noise scenarios illustrated in Fig. 1e, where curvature estimates reflected noise-driven structure rather than the underlying response geometry. In contrast, indirect curvature estimates were substantially lower (all below 100°) and revealed a more regular pattern, with later areas exhibiting progressively straighter trajectories (Fig. 2a, black symbols; AC: 96.46°, sPMv: 97.00 °, AG: 94.71°, PFC: 92.72°). This pattern is consistent across datasets. We analyzed data from 9 subjects who listened to up to 3 stories, each repeated at least 5 times, yielding 17 unique datasets (each shown as a thin line in Fig. 2b).

**Figure 2.**
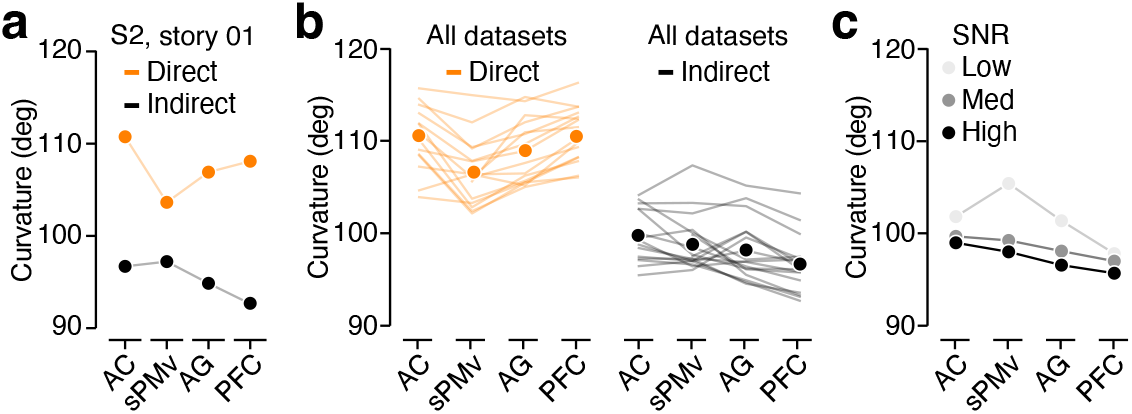
Indirect neural curvature estimates support the temporal straightening hypothesis. (**a**) fMRI response curvature estimates obtained through two estimation methods (orange: direct vs black: indirect) across four different brain regions for an example subject (S2, listening to story 01). The direct method yields consistently higher curvature estimates and does not show a systematic hierarchical organization. The indirect method yields lower curvature estimates and shows a progressive decrease in curvature from AC to PFC. (**b**) Curvature estimates for all datasets (each line shows one dataset; symbols show the average). (Left) Direct method. (Right) Indirect method. (**c**) Indirect curvature estimates computed separately for voxels with low, medium, and high SNR. Curvature decreases from AC to PFC within each SNR bin, indicating that the hierarchical decrease in curvature is not driven by differences in SNR

To test the temporal straightening hypothesis, we first compared curvature between the lowest (AC) and highest (PFC) levels of the processing hierarchy. A systematic difference was evident in the indirect estimates (median difference = −3.301°, *W* = 6, *P <* 0.001, Wilcoxon signed-rank test), but not in the direct estimates (median difference = −0.188°, *W* = 56, *P* = 0.847). We complemented this simple analysis with a more complex one (a linear mixed-effect model) that included all four brain areas (treated as an ordered numeric predictor) and allowed for subject- and story-specific effects. This confirmed that indirect curvature estimates decrease along the hierarchy (slope = −1.002 ± 0.179 SE, *P <* 0.001; Fig. 2b, right), whereas direct curvature estimates do not exhibit this trend (slope = 0.318 ± 0.262 SE, *P* = 0.225; Fig. 2b, left). We verified that these results cannot be explained by differences in ROI size (Extended Data Fig. 2). Finally, we asked whether cross-area differences in fMRI signal quality might partially drive these effects. We estimated the signal-to-noise ratio (SNR) from voxel-averaged responses within each ROI for every subject and story and found that SNR did not exhibit a systematic hierarchical trend. This suggests that cross-area SNR differences cannot explain our main results. To test this directly, we binned all voxels by SNR-level (three groups) and repeated the indirect curvature analysis for each of the voxels in each ROI and each bin. Within each SNR level, curvature decreases along the hierarchy (lowest SNR bin: slope = −1.849 ± 0.442 SE, *P <* 0.001; middle bin: slope = −0.938 ± 0.194 SE, *P <* 0.001; highest bin: slope = −0.978 ± 0.220 SE, *P <* 0.001; Fig. 2c). We conclude that response trajectories robustly straighten along the language processing hierarchy.

### A brain-wide map of trajectory curvature

The connection between representational timescale and trajectory curvature can be exploited to estimate implied population trajectory curvature from single voxel responses. To complement the ROI analyses, which pool responses across voxels to maximize SNR, we next examined curvature on a voxel-by-voxel basis. While sacrificing some noise robustness, a single-voxel approach allows us to probe finer-grained spatial structure. This provides an opportunity to test whether hierarchical trends are also present within individual ROIs, as we would expect along the medial-lateral axis in auditory cortex [11]. We computed voxelwise implied trajectory curvature maps by estimating AR(1) coefficients separately for each voxel across the whole brain. To ensure reliable estimates, we filtered voxels based on trial-wise coherence, excluding those with low response reliability. In an example subject (Fig. 3), curvature values were highest in medial auditory cortex and decreased progressively both within AC (from primary auditory cortex, A1, to more lateral regions) and along the broader hierarchy toward higher-order speech-processing areas (more examples shown in Extended Data Fig. 3). The same gradient was evident in the group-average map (Fig. 4). Specifically, curvature was highest in auditory cortex and smallest in the angular gyrus, superior precuneus and anterior PFC. Within-area gradients were evident in the auditory cortex (primary to lateral regions), the precuneus (inferior to superior), and PFC (posterior to anterior). This topographic structure resembles the large-scale organization of connectivity patterns [37] and local synaptic strength [38]. We conclude that hierarchical straightening in the speech processing system reflects a continuous cortical motif, and that it may be linked to other large scale organizing principles.

**Figure 3.**
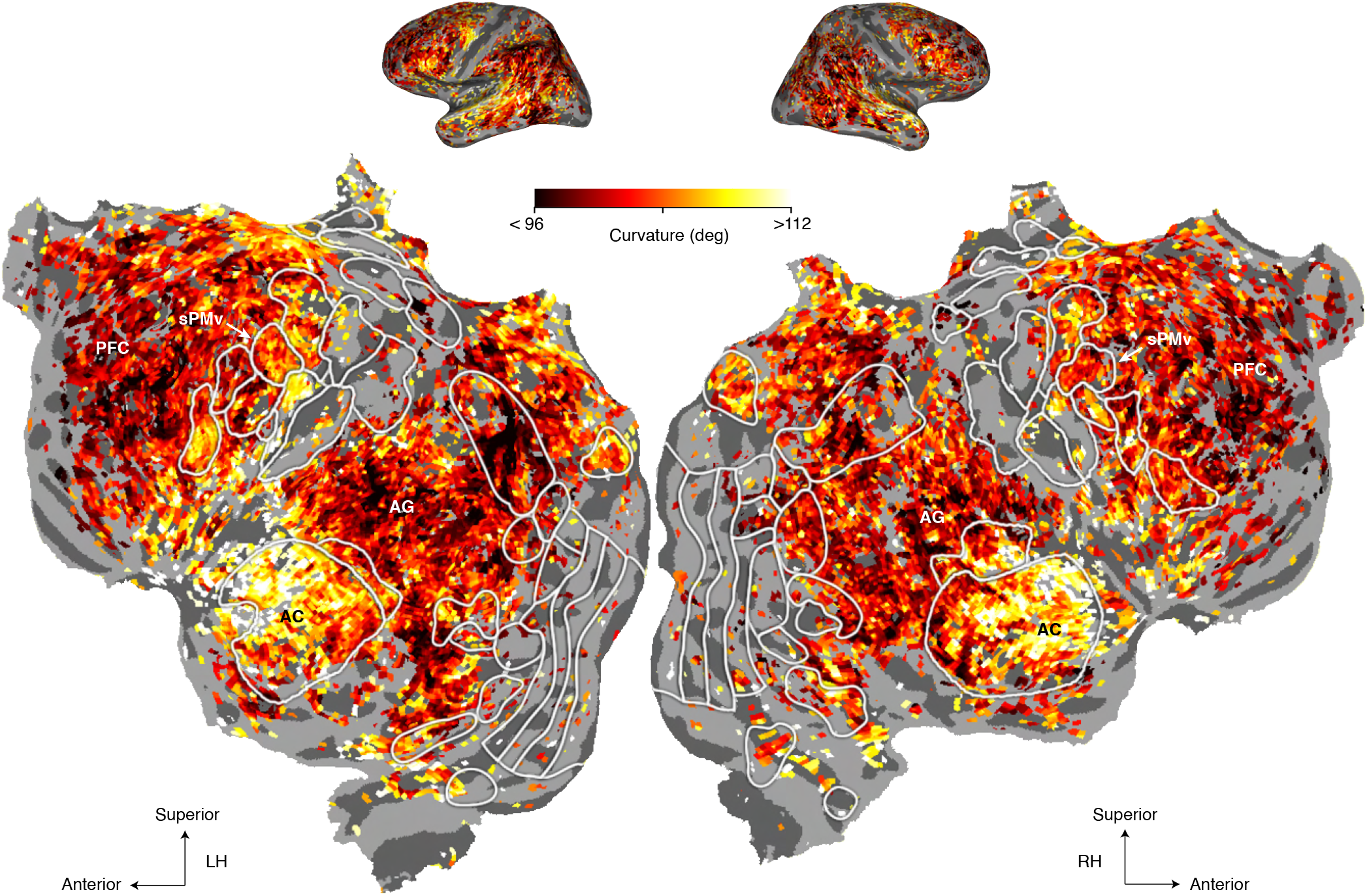
Subject-level cortical maps of implied trajectory curvature. Curvature estimates are shown separately for each subject while listening to story 01, using the same indirect estimation procedure as in Fig. 3. For each voxel, we fit an AR(1) model to the run-averaged response time course and converted the estimated autoregressive coefficient to implied trajectory curvature using the analytic relationship between *ϕ* and *θ*. Only voxels with coherence values above the 80th percentile and greater than 0.05 are shown. Across subjects, implied curvature is highest in medial AC and decreases progressively both within AC and along the broader cortical hierarchy toward higher-order speech processing areas. These maps show that the spatial gradient observed in the example subject is consistent across individual participants.

**Figure 4.**
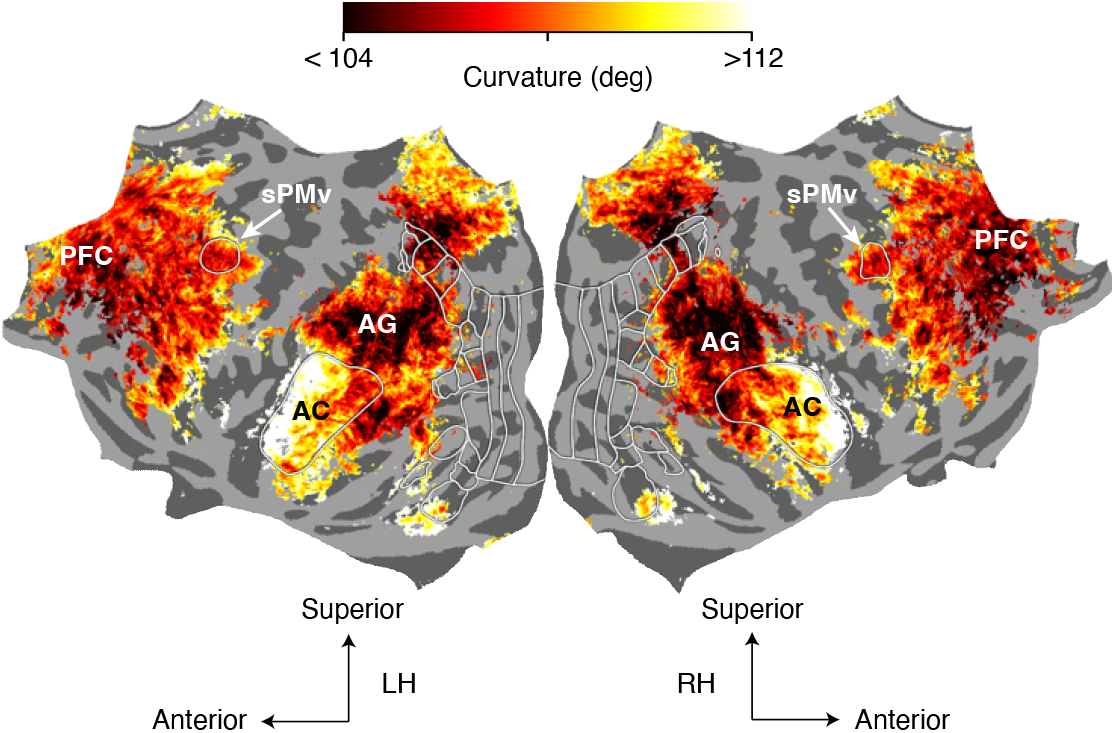
Average cortical map of implied trajectory curvature across all datasets and subjects. For each subject and story, voxel-wise AR(1) coefficients were estimated from the run-averaged response time course, projected onto the cortical surface, and aligned to the fsaverage template. AR(1) maps were then averaged across subject–story datasets and converted to implied trajectory curvature using the analytic relationship between *ϕ* and *θ*. Coherence was computed voxel-wise and averaged in the same fsaverage space; only vertices with mean coherence above the 60th percentile and greater than 0.05 are shown. The group-average map preserves the spatial organization observed in the example subject: inferred curvature is highest in AC and decreases along the cortical hierarchy toward higher-order speech processing regions. This pattern indicates that the cortical gradient in trajectory curvature is reproducible across subjects and stories.

### Temporal straightening is tailored to the statistics of natural speech

We hypothesize that the progressive straightening of response trajectories along the cortical speech processing hierarchy arises from a sequence of transformations adapted to the statistics of natural speech. If so, straightening effects should only emerge in systems exposed to speech. Moreover, these effects should be weaker or absent for auditory inputs whose statistical properties deviate from those of speech. To test these hypotheses, we analyzed responses of an artificial speech processing system. We focused on WavLM, a deep neural network trained on large-scale natural speech data [39] (Fig. 5a). Models of this class have been shown to resemble cortical speech processing: voxel activity in earlier regions of the cortical language hierarchy, which primarily encode acoustic features, is best predicted by early layers of the model, whereas voxel activity in later language regions, which encode phonetic and semantic information, is best predicted by upper-middle layers [34].

**Figure 5.**
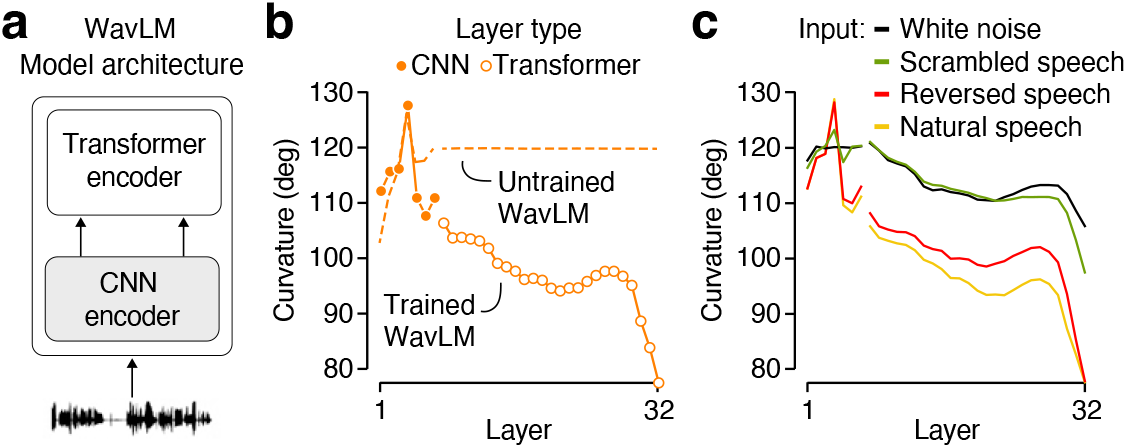
Temporal straightening in WavLM is tailored to the statistics of natural speech. (**a**) Schematic of the WavLM architecture. Auditory speech stimuli are first processed by a 7-layer CNN encoder, which provides input to a 25-layer transformer encoder. (**b**) Layer-wise trajectory curvature estimated using the direct estimation procedure from model hidden-state representations elicited by the same auditory speech stimuli used in our fMRI experiments. Closed symbols: CNN layers; open symbols: Transformer layers. Dotted lines show the same analysis performed in the untrained model. Progressive straightening emerges following training on natural speech data. (**c**) Hierarchical evolution of WavLM trajectory curvature for different input stimuli. We probed the network with natural speech (yellow), reversed speech (red), phase-scrambled speech (green), and white noise (black). Temporal straightening was strongest for natural speech, reduced for reversed speech, and substantially attenuated for phase-scrambled speech and white noise.

We first presented the same natural speech stimuli used in the fMRI experiments to WavLM and computed trajectory curvature for all layers of the network. We used the direct estimation procedure, which for this noise-free system captures the ground truth, and analyzed the model before and after training on speech data. Along the network hierarchy, trajectory curvature decreases by roughly 40° in the trained model (Fig. 5b, orange symbols, solid line). In contrast, no such effect was evident in the untrained version of the same architecture (Fig. 5b, dashed line). Temporal straightening is thus not a trivial consequence of the model architecture but instead emerges through training on natural speech.

To test the specificity of temporal straightening effects, we created several perturbed versions of the natural speech stimuli used in the fMRI experiments. These perturbations differed in how much speech-like structure they preserved. First, we created reversed speech, which is different from natural speech semantically, but preserves many local acoustic features of the original signal, including spectrotemporal patterns, phonetic-like transitions, amplitude modulations, and correlations across time and frequency, just in reverse order. We still observe progressive straightening across layers of the network; however, the magnitude of straightening in middle layers was reduced compared to natural speech (Fig. 5c, red vs yellow). Scrambled speech provides a stronger perturbation: it preserves the amplitude spectrum of the original waveform but disrupts the phase relationships that organize speech over time. This manipulation substantially reduced temporal straightening (Fig. 5c, green vs yellow). White noise, which lacks both the speech-like amplitude spectrum and the phase-dependent temporal organization, produced the smallest straightening effects (Fig. 5c, black). Together, these in silico experiments suggest that temporal straightening in speech processing systems arises from a sequence of transformations tailored to the specific statistics of speech.

### Temporal straightening generalizes across temporal resolutions

Our WavLM and fMRI analyses differ in two important regards: curvature can be estimated directly from WavLM representations whereas fMRI requires an indirect estimate, and the two systems time-varying responses have very different temporal resolutions. To connect both sets of analyses, we first compared direct and indirect curvature estimates in WavLM using the natural speech stimuli shared with the fMRI experiment (Methods). The indirect procedure captures the progressive decrease in trajectory curvature along the hierarchy, but underestimates the overall magnitude of this decrease (Fig. 6a, black vs orange). Thus, to the extent that the same bias applies to fMRI, the resulting estimates likely provide a conservative measure of the true effect size of temporal straightening in the human brain.

**Figure 6.**
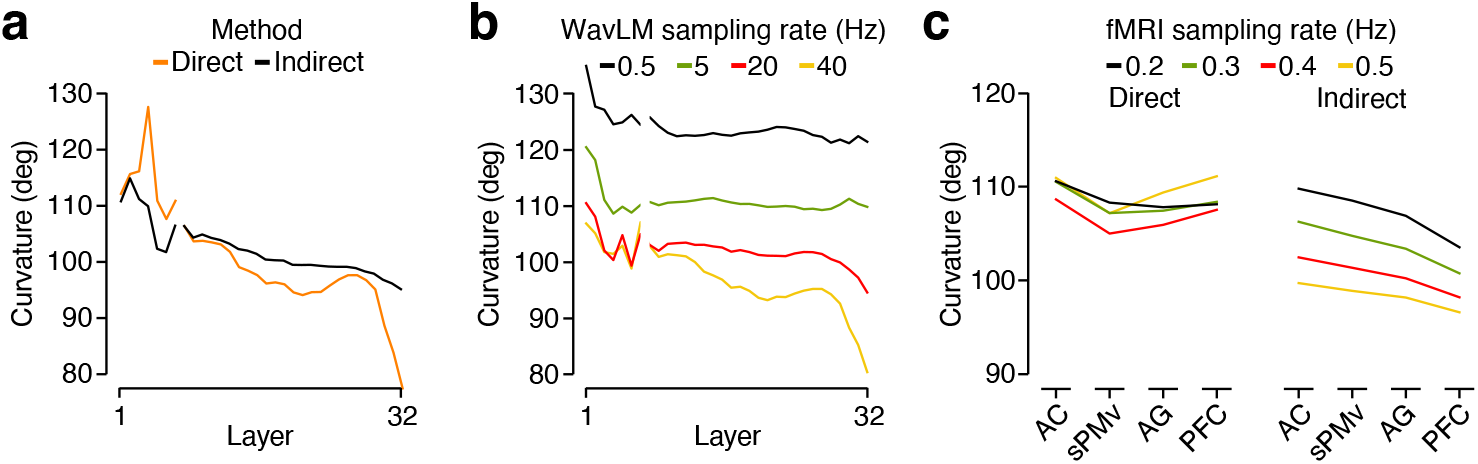
Temporal straightening generalizes across temporal resolutions. (**a**) Comparison of direct (orange) and indirect (black) trajectory curvature estimates in WavLM responses to natural speech. Because the network is noiseless, the direct method provides a ground-truth estimate of trajectory curvature. The indirect method captures the overall decreasing trend along the network hierarchy, but underestimates the magnitude of this effect. (**b**) WavLM trajectory curvature computed from network responses resampled to different sampling rates. Hierarchical straightening is evident at each temporal resolution. Overall curvature was higher at slower sampling rates, whereas the decrease in curvature across the network hierarchy was larger at faster sampling rates. (**c**) fMRI trajectory curvature computed from brain responses resampled to different sampling rates. Hierarchical straightening is evident at each temporal resolution for the indirect estimates, but not for the direct estimates. As in WavLM, overall curvature was higher at slower sampling rates, whereas the decrease in curvature across the cortical hierarchy was larger at faster sampling rates. Lines show the average curvature estimates across all datasets. Left: direct method. Right: indirect method.

Speech contains structure at slow as well as fast temporal resolutions [40], all of which can be exploited for prediction. This raises the question of whether temporal straightening occurs across multiple temporal resolutions. To address this question, we progressively downsampled WavLM activations and computed trajectory curvature at each sampling rate, using the direct estimation procedure. Across all layers of the network, slower sampling rates were associated with higher curvature (Fig. 6b). Because this offset was already evident in the initial encoding layer, we interpret differences in absolute curvature across sampling rates as reflecting differences in the temporal structure available in the input. Our primary question, however, concerned the relative change in curvature across the processing hierarchy. Critically, hierarchical temporal straightening was evident at every sampling rate, including the fMRI sampling rate of 0.5 Hz (Fig. 6b, black line). This pattern suggests that temporal straightening may be a representational motif across multiple temporal resolutions in the human speech processing system. To test this possibility, we analyzed the fMRI responses at temporal resolutions even slower than the native sampling rate, using both the direct and indirect estimation procedures. As before, the direct estimates did not exhibit obvious structure (Fig. 6c, left). In contrast, the indirect estimates exhibited the same qualitative pattern as the downsampled WavLM trajectories (Fig. 6c, right). Because different downsampling methods alter the source signal in different ways, we repeated these analyses using an alternative downsampling procedure and observed the same pattern of results (Extended Data Fig. 4). We conclude that temporal straightening occurs across multiple temporal resolutions in WavLM and the human speech processing system.

### Curvature and timescale in populations of spiking neurons

The structure of fMRI brain responses only partly reflects patterns of spiking activity in large neural populations [41]. We therefore asked whether the link between the representational timescale of single-unit activity and the curvature of population response trajectories extends to networks of spiking neurons. To address this question, we leveraged the statistical framework of *continuous modulated Poisson processes* [42, 43] and simulated spiking activity for populations of neurons driven by time-varying inputs. This framework provides an accurate account of cortical neuron response statistics [44, 45, 46, 42] and offers simple parametric control over the timescale of neural signals and neural noise (Methods; Fig. 7a). We analyzed spiking activity for simulated populations of 100 neurons, with all neurons in a population sharing the same signal and noise timescales, while the exact temporal profile of each component varied across neurons (examples shown in Fig. 7b,c). For each population, we computed trajectory curvature in a representational distance space, yielding a metric conceptually analogous to the curvature measures used for fMRI and WavLM (Methods).

**Figure 7.**
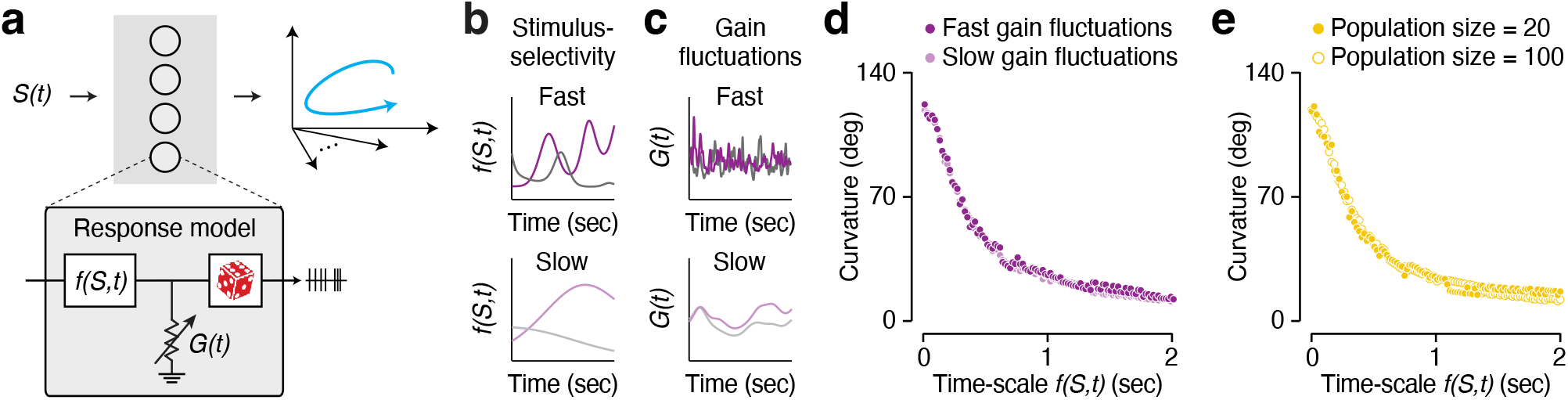
Curvature and timescale in simulated populations of spiking neurons. (**a**) We used the continuous modulated Poisson model [42] to simulate spiking activity for populations of neurons driven by time-varying inputs. The pattern of spiking activity is structured by each neuron’s stimulus selectivity (f(S,t)), time-varying excitability (G(t)), and Poisson noise (dice). The model’s closed form expressions for spike count distributions enabled us to express population trajectories in a representational distance space. Trajectory curvature was computed in this space. (**b**) Example units (two per panel) with fast vs slow signal timescale (top vs bottom). (**c**) Example units (two per panel) with fast vs slow noise timescale (top vs bottom). (**d**) Neural trajectory curvature plotted as a function of signal timescale. Each symbol illustrates the curvature of a single population trajectory (population size = 100 units; counting window = 200 ms), color indicates noise timescale (fast vs slow). Fast signal timescales yielded trajectories with high curvature, whereas slow signal timescales yielded nearly straight trajectories. The timescale of neural noise had no effect on trajectory curvature. (**e**) Neural trajectory curvature plotted as a function of signal timescale for populations composed of 20 (open symbols) vs 100 (closed symbols) units. Plotting conventions are identical to panel d. Trajectory curvature depended primarily on representational timescale and was unaffected by population size.

We considered populations with varying representational timescales and found a prominent relationship between the timescale of neural signals and trajectory curvature (Fig. 7d). Fast timescales (on the order of 10 ms) yielded trajectories with curvature exceeding 100°. Slow timescales (on the order of 1 s) yielded much straighter trajectories with curvature approaching 0°. In contrast, the timescale of neural noise had little effect on trajectory curvature (Fig. 7d, dark vs light symbols). For AR(1) processes, trajectory curvature is determined by representational timescale and is independent of population size. To test whether this property also holds in networks of spiking neurons, we compared populations composed of 100 vs 20 neurons. Again, curvature varied with representational timescale but not with the number of neurons in the population (Fig. 7e, open vs closed symbols). Together, these simulations suggest that the link between timescale and curvature is a robust property of neural population codes, extending beyond autoregressive models and artificial neural networks.

## Discussion

In this work, we suggest that the human speech processing system exhibits temporal straightening as a representational motif, with trajectory curvature progressively decreasing along the cortical hierarchy. We further propose that this geometric transformation is governed by the changing temporal dynamics of neural responses and establish a principled relationship between representational timescale and the curvature of population trajectories. We first identified this relationship in simple autoregressive models, where it can be derived analytically, and found that it generalizes to deep neural network representations (WavLM), as well as to models of spiking activity in neural populations. Across these systems, slower temporal dynamics consistently give rise to straighter population trajectories. We leveraged this relationship to develop an indirect approach to estimate curvature from fMRI data that mitigates the biasing effects of measurement noise. Together, these results link the goal of temporal prediction to representational timescale and the geometry of population trajectories.

Our findings offer a perspective on language processing in which the statistical structure of natural speech is reflected in the geometry of neural representations. This perspective differs from coding schemes built on the explicit representation of prediction errors [15, 16, 17, 18]. Instead, we propose that predictable speech transitions manifest as smoothly evolving neural population trajectories. The advantage of this coding scheme is that prediction does not require a separate, resource-intensive feedback process; instead, predictions are implicitly embedded in the structure of the representation itself. This allows the brain to anticipate future representations efficiently without constructing explicit predictions, and to flexibly support multiple aspects of speech processing using the same underlying representation. For example, when speech is noisy or ambiguous, constraining the evolving trajectory along a straighter path effectively leverages prior knowledge. Moreover, local departures from smooth trajectories could provide a geometric signature of surprisal.

The absolute magnitude of the straightening effects observed in the fMRI data appears modest (∼ 5–15°). This is in part due to a downside of our indirect approach for estimating trajectory curvature: its limited dynamic range. Real trajectories can have curvatures ranging from 0° to 180°, but AR(1) processes can only exhibit curvatures between 90° and 180°. A second reason is that fMRI responses have a coarse spatiotemporal resolution compared to the neural events of interest. Analyses of WavLM representations provide an informative comparison because they do not suffer from the spatiotemporal limitations of fMRI. In this noiseless system, we can directly measure trajectory curvature and find that at the fastest temporal resolution, straightening effects are substantially stronger (∼ 40°) than our fMRI estimates. Moreover, the continuous modulated Poisson model indicates that much larger curvature ranges are possible in neural populations. Varying the timescale of stimulus-selective responses over biologically plausible ranges (milliseconds to seconds), yields trajectories whose curvature spans a range up to ∼ 120°. Together, these findings suggest that the fMRI straightening effects likely reflect an underestimate of the underlying neural effects. Future work using intracranial recordings of spiking activity during naturalistic speech processing will be critical for directly characterizing population trajectories and testing the magnitude of temporal straightening across the cortical hierarchy.

The brain supports perception, cognition and action through the collective activity of large populations of neurons. Two key aspects of this activity are its dynamics and its geometry. The dynamics provide insight into the computational strategies employed by hard-wired neural networks [47], while the geometry determines the resulting perceptual, cognitive, and motor capabilities [48]. Although both aspects have mostly been studied in isolation, many recent investigations of spiking activity have investigated their relation. These efforts have clarified the neural basis of time-perception [49], decision-making [50], flexible inference [51], compositional cognition [52], and motor planning [53]. Here, we have shown that one of the simplest characteristics of response dynamics (representational timescale) is related to one of the most basic geometric aspects of a time-varying representation (trajectory curvature). This insight is not only valid at the level of spiking activity but was also useful to analyze time-varying fMRI responses. Our study thus extends the endeavor to link neural dynamics and geometry to non-invasive recording methods. Importantly, however, this link is computational rather than mechanistic. Our results do not specify the circuit-level mechanisms that give rise to different representational timescales in different brain regions. In both the visual and speech processing system, representational timescales increase along the cortical hierarchy, reflected in progressively longer temporal receptive windows [54, 55, 31, 33, 56]. This trend may arise from gradual changes in synaptic strength [38], cellular dynamics, recurrent connectivity, or cross-area interactions [37]. Determining the mechanistic basis of the geometric transformations we observed will be an important direction for future work.

Our findings extend recent discoveries of temporal straightening in the primate visual system [19, 28], in hierarchical models for video prediction [57], in world models that predict how environments evolve under actions [21, 58], and in LLMs trained on next-word prediction [25, 59]. This commonality is surprising, given the stark structural and functional differences across these systems. Why do all these systems favor straighter representations over alternative geometric structures? One possibility is that straightening is imposed by the basic architecture of the system. Hierarchical networks may automatically reshape raw inputs through pooling operations, transforming high-dimensional curved input trajectories into low-dimensional smoother representations which coincidentally are easier to extrapolate. This seems unlikely, given that straightening does not occur in untrained speech representation models (Fig. 5b) [25]. Network architecture is thus not a sufficient explanation for temporal straightening. A more likely alternative is that the computational objective of prediction exposes the underlying high-level temporal structure of the natural sensory environment. Natural signals such as speech and video are not random, but arise from sources that are structured in time, even if this is not evident in the resulting raw sensory inputs. Specifically, light is reflected by matter that coheres into stable structures through fundamental physical forces while the acoustic events caused by speech follow a hierarchical temporal organization that includes rhythmic, nested units and prosodic features. The progressive increase of temporal representational regularity along a network hierarchy may thus reflect the recovery of these more stable latent high-level features embedded in the sensory stream. Consistent with this, a recent study found that neural networks that used temporal straightening as a learning principle for representing videos contained predictive neural embeddings that also capture and separate geometric, photometric, and semantic object features [60]. Out of all possible regular trajectories, straight ones have the specific benefit that representational distance systematically grows with separation in time, facilitating discrimination and identification of distinct temporal events. While this account is somewhat speculative, the accumulating evidence for temporal straightening across diverse biological and artificial systems suggests that it reflects a general principle by which hierarchical systems organize their internal dynamics to make the future more predictable.

## METHODS

### Modeling: Relationship between AR(1) dynamics and trajectory curvature

We sought to analytically relate the curvature of population response trajectories to the temporal dynamics of an underlying autoregressive process. Consider an N-dimensional response vector *y*_*t*_ ∈ R^*N*^ denote the population response at time *t*, evolving according to an autoregressive process of order one (AR(1)):

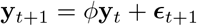

where 0 *< ϕ <* 1 and ***ϵ***_*t*_ ∼ *N* (0, *σ*^2^*I*_*N*_). *I*_*N*_ denotes the identity matrix. Under this model, each dimension evolves independently and the process is stationary with covariance

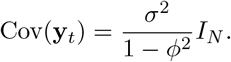

To quantify curvature, we consider successive displacement vectors in population space:

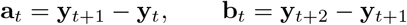

and define the instantaneous curvature angle as

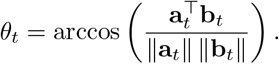

Using the AR(1) dynamics, the displacement vectors can be written as

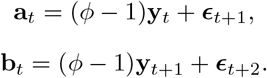

Substituting **y**_*t*+1_ = *ϕ***y**_*t*_ + ***ϵ***_*t*+1_ into **b**_*t*_ yields

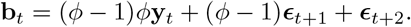

Since **y**_*t*_ is independent of future innovations and the innovations are mutually independent, we obtain

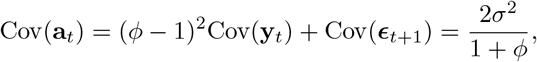

and similarly

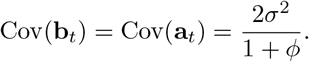

The covariance between successive displacements is

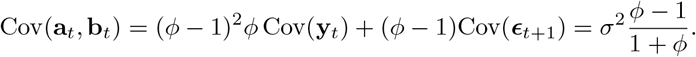

It follows that the correlation between successive displacement vectors is

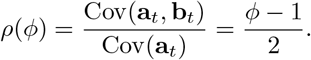

The vectors **a**_*t*_ and **b**_*t*_ are jointly Gaussian and isotropic across dimensions, with covariance

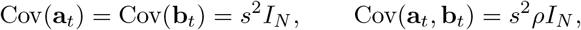

for some scalar *s*^2^. The curvature depends only on the normalized inner product

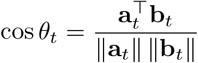

which is invariant to scale.

For isotropic jointly Gaussian vectors with correlation *ρ*, the expected angle between the vectors is

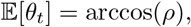

independently of the dimensionality *N* ≥ 2.

Substituting *ρ* yields

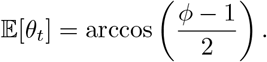

### Modeling: Evaluating direct and indirect curvature estimates in AR(1) simulations

We simulated population time series using a first-order autoregressive (AR(1)) process to examine how noise affects direct and indirect estimates of trajectory curvature. For each simulation, we generated responses for *N* = 100 voxels over *T* = 241 time points, across a range of autoregressive coefficients (*ϕ* ∈ [0.1, 1], 10 values) and bootstrap repetitions (500 iterations). To simulate repeated measurements, we generated 10 runs per condition. For each voxel *v*, the latent time series evolved according to

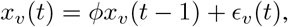

where *ϵ*_*v*_(*t*) is a zero-mean innovation term with total variance *σ*^2^. In the noiseless condition, the innovation term was decomposed into a private component and a shared component across voxels, such that

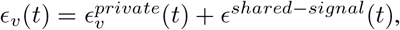

where the total variance *σ*^2^ was partitioned such that a fraction 1 − *α* was private to each voxel and a fraction *α* was shared across voxels. Both components were independent across time, and only the private component was independent across voxels.

In noisy simulations, the shared component was further decomposed into signal and noise components, such that

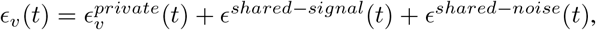

where the shared variance *ασ*^2^ was divided into a signal component that was identical across voxels and runs, with variance *αβσ*^2^, and a noise component that varied across runs, with variance *α*(1 − *β*)*σ*^2^. The same value of *α* was used as in the noiseless condition. For each run, a new realization of the shared noise component was drawn, while private and shared signal components were held fixed across runs. After generating the AR(1) time series, additive Gaussian measurement noise was applied independently to each voxel and time point, with standard deviation *σ*_*meas*_ = 1.5 (low noise) or 3.0 (high noise). Responses were then averaged across runs to mimic trial averaging in fMRI.

Direct curvature was computed from population trajectories obtained by averaging simulated responses across 10 runs, by treating each time point as a point in voxel space. Step vectors were defined as **v**(*t*) = **x**(*t* + 1) − **x**(*t*), where **x**(*t*) denotes the population response across voxels at time t. The angle between successive steps was computed as

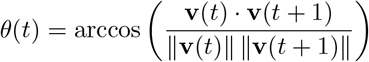

and expressed in degrees. Direct curvature was defined as the average of *θ*(*t*) across time. Time points for which step magnitudes were numerically close to zero were excluded to avoid instability in the angle computation. We first simulated noiseless latent trajectories with different ground-truth curvature by varying the autoregressive coefficient *ϕ*. Ground-truth curvature was computed using the analytical relationship derived above, (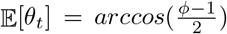). In noisy simulations, shared variability across voxels and additive measurement noise (low and high) were introduced as described above, and direct curvature was recomputed on the resulting population trajectories. Error bars indicate the 95% confidence intervals obtained from the bootstrapsamples.

Indirect curvature was estimated by fitting an AR(1) model to the averaged population trajectories (across 10 runs) and extracting the autoregressive coefficient *ϕ*. Curvature was then computed from the estimated *ϕ* using the same analytical relationship. This procedure was applied to both noiseless and noisy simulations. Error bars indicate 95% confidence intervals obtained from bootstrap samples.

### MRI data

#### subjects and stimuli

We analyzed functional magnetic resonance imaging (fMRI) data from 9 human subjects as they listened to English language podcast stories over Sensimetrics S14 headphones. Data from seven subjects (S1–S7) were obtained from previously published, publicly available datasets [30]. In addition, we included data from two new subjects (BFD001, BFD003) collected using the same experimental paradigm and acquisition procedures. Subjects were not asked to make responses, but simply listened attentively to the stories. We restricted our analyses to stories with repeated presentations. Three stories from the podcast *The Moth Radio Hour* were included: *Where There’s Smoke, From Boyhood to Fatherhood*, and *On Approach to Pluto*. Each 10-15 minute story was played during a separate scan. Subjects S1–S3 listened to all three stories (10, 6, and 4–5 repetitions, respectively). The remaining subjects (S4–S7, BFD001, BFD003) contributed data only for *Where There’s Smoke*, with 5–9 repetitions per subject. Across all subjects and stories, this yielded a total of 15 subject-story datasets with at least four repetitions each. For all analyses, responses were averaged across repetitions. Only voxels within 8 mm of the mid-cortical surface were analyzed, yielding roughly 90,000 voxels per subject.

#### MRI acquisition

Details of the MRI methods can be found in the original publications [30], but important points are summarized here. MRI data were collected on a 3T Siemens Skyra scanner at University of Texas at Austin using a 64 channel Siemens volume coil. Functional scans were collected using a gradient echo EPI sequence with repetition time (TR) = 2.00 s, echo time (TE) = 30.8 ms, flip angle = 71°, multi-band factor (simultaneous multi-slice) = 2, voxel size = 2.6*mm*×2.6*mm*×2.6*mm* (slice thickness = 2.6*mm*), size = × 84, and field of view = 220*mm*. Anatomical data were collected using a T1-weighted multi-echo MP-RAGE sequence with voxel size = 1*mm* × 1*mm* × 1*mm* following the Freesurfer morphometry protocol [61].

All subjects were healthy and had normal hearing. The experimental protocol was approved by the Institutional Review Board at the University of Texas at Austin. Written informed consent was obtained from all subjects.

#### Preprocessing

All functional data were motion corrected using the FMRIB Linear Image Registration Tool (FLIRT) from FSL 5.0. FLIRT was used to align all data to a template that was made from the average across the first functional run in the first story session for each subject. These automatic alignments were manually checked for accuracy. Low frequency voxel response drift was identified using a 2nd order Savitzky-Golay filter with a 120 second window and then subtracted from the signal. To avoid onset artifacts and poor detrending performance near each end of the scan, responses were trimmed by removing 20 seconds (10 volumes) at the beginning and end of each scan, which removed the 10-second silent period and the first and last 10 seconds of each story. The mean response for each voxel was subtracted and the remaining response was scaled to have unit variance.

#### Defining regions of interest (ROIs)

Regions of interest (ROIs) were localized separately for each subject. The anatomical ROIs, angular gyrus (AG) and prefrontal cortex (PFC) were defined using Freesurfer labels. AG was defined using the ‘inferiorparietal’ label, and PFC was defined using the ‘parsopercularis’, ‘parstriangularis’, ‘superiorfrontal’, ‘rostralmiddlefrontal’, ‘caudalmiddlefrontal’, ‘frontalpole’ labels. Functional ROIs were defined using two localizer tasks: an auditory cortex localizer and a motor localizer. Auditory cortex localizer data were collected in one 10-minute scan. The subject listened to 10 repeats of a 1-minute auditory stimulus containing 20 seconds each of music (Arcade Fire), speech (Ira Glass, This American Life), and natural sound (a babbling brook). To determine whether a voxel was responsive to auditory stimulus, the repeatability of the voxel response across the 10 repeats was calculated using an F-statistic. This map was used to define the auditory cortex (AC). Motor localizer data were collected in two identical 10-minute scans. The subject was cued to perform six different tasks (“hand”, “foot”, “mouth”, “speak”, “saccade”, and “rest”) in a random order in 20-second blocks. For the “speak” cue, subjects were instructed to self-generate a narrative without vocalization. The weight map for the “speak” cue was used to define the superior ventral premotor speech area (sPMv).

### fMRI analysis

#### Direct and indirect method

We applied both direct and indirect procedures to estimate the curvature of population trajectories from fMRI responses across cortical regions. For each dataset (one subject, one story), the direct method was applied to trial-averaged voxel responses. Within each ROI, population activity was treated as a trajectory in a high-dimensional space, with each voxel representing a dimension. Trajectory curvature was then computed as the average angle between consecutive time points (TRs), using the same definition described above. For the indirect method, responses were first averaged across voxels within each ROI, in addition to trial averaging, to improve SNR. An AR(1) model was then fit to the averaged response to estimate the autoregressive coefficient *ϕ*, from which curvature was computed using the analytical relationship described above.

#### Statistical analysis

Differences in curvature between the lowest (AC) and highest (PFC) levels of the hierarchy were assessed using a two-sided Wilcoxon signed-rank test across datasets (one subject–story pair). To evaluate trends across the full hierarchy, we fit a linear mixed-effects model with curvature as the dependent variable and ROI position treated as an ordered predictor, with subject and story included as random effects.

#### Control analysis for SNR

To assess whether differences in SNR across cortical regions could influence curvature estimates, we performed a control analysis in which comparisons across ROIs were made at matched SNR levels. For each voxel, SNR was estimated from repeated presentations of the same stimulus by decomposing responses into signal and noise components. The signal was defined as the mean response across repetitions, and noise as the deviation of each repetition from this mean. SNR was computed as the ratio of signal variance to total variance. Within each ROI, voxels were grouped into three SNR bins based on these estimates, using bin edges defined globally across all ROIs. For each bin, responses were averaged across repetitions and then across voxels, yielding a single time course per bin for each dataset (subject-story pair). The indirect procedure was then applied to these bin-wise averaged responses: AR(1) models were fit to estimate the autoregressive coefficient *ϕ*, and curvature was computed from *ϕ* using the analytical relationship described above. The same statistical tests were then applied to the bin-wise curvature estimates.

### WavLM experiments

In this work, we used WavLM, a self-supervised speech model trained to learn representations from raw speech waveforms. WavLM is based on the HuBERT framework and combines masked speech prediction with architectural and training modifications designed to improve both spoken-content modeling and speaker-related representations [39]. In the main analysis, we used the pre-trained Large variant from Hugging Face. This model contains a convolutional feature encoder followed by a Transformer contextualizer and was pretrained on 16-KHz English speech from large-scale unlabeled speech corpora.

#### Direct and indirect curvature estimation procedures

For both estimation procedures, audio stimuli were first resampled to 16 kHz and passed through the frozen model without fine-tuning. We extracted layer-wise hidden-state representations from the network. For a given layer, the activation vector at each time point defined a point in representational space, with each dimension corresponding to the response of one unit. The sequence of activation vectors over time therefore defined a trajectory through representational space. The direct method estimated curvature by computing the unsigned angle between consecutive segments of this trajectory, and layer-wise curvature was defined as the average angle across time. Indirect curvature estimates were obtained by fitting an AR(1) model to each unit’s response time series. For each layer, we averaged the resulting autoregressive coefficients across units and converted this average coefficient into a curvature estimate using the analytic relationship between the AR(1) coefficient and expected curvature (*θ* = arccos((*ϕ* − 1)*/*2)).

#### Perturbed natural speech stimuli

We made several perturbed versions of the natural speech stimuli, passed them through the pre-trained model and extracted layer-wise hidden-state representations as described above. Reversed speech was generated by reversing the temporal order of the natural speech waveform. For the phase-scrambled speech stimulus, we first performed a Fourier transform on the natural speech waveform to obtain its amplitude and phase spectra. We kept the amplitude spectrum intact and replaced the original phase spectrum with random phases sampled uniformly from 0 to 2*π*. We then transformed the resulting spectrum back into the time domain. The white noise stimulus was generated by sampling independent Gaussian values at each time point. All perturbed stimuli were scaled to match the RMS amplitude of the original speech waveform.

#### Downsampling WavLM activations and fMRI data

To examine temporal straightening across different temporal resolutions, we first downsampled the layer-wise WavLM activation time courses directly. For each layer, activations were resampled from their native sampling rate to a set of lower sampling rates (40 Hz, 20 Hz, 5 Hz, and 0.5 Hz) using polyphase resampling. The main analysis used direct resampling without prior low-pass filtering. Trajectory curvature was then computed from the downsampled activation time courses using the direct estimation procedure described above.

For the fMRI data, we applied the same downsampling procedure to the run-averaged voxel time courses separately for each dataset (one subject-story pair) and then averaged across datasets for each ROI and temporal resolution. For the direct estimation procedure, voxel time courses were downsampled to lower sampling rates (0.4 Hz, 0.3 Hz, and 0.2 Hz), and population trajectory curvature was computed across all voxels within each ROI. For the indirect estimation procedure, we first downsampled the run-averaged voxel time courses and then averaged the downsampled time courses across voxels within each ROI. We fit an AR(1) model to the resulting ROI-averaged time course to estimate the autoregressive coefficient, which was converted into a curvature estimate using the analytic relationship derived above.

As a control, we repeated these analyses after applying an anti-aliasing low-pass filter before resampling. Specifically, we used a fourth-order Butterworth low-pass filter with a cutoff frequency set to 80% of the target Nyquist frequency, applied with zero-phase filtering. These results are shown in Extended Data Figure 4.

### Populations of spiking neurons

To relate the timescale of single-unit spiking activity to the curvature of population response trajectories, we leveraged the *continuous modulated Poisson* (CMP) framework [42]. Under this model, the time-varying tuning function *f*_*i*_(*S, t*) of neuron *i* = 1, …, *N* in response to a stimulus *S* is governed by an exponentiated Gaussian process:

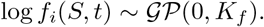

Here, *K*_*f*_ is a radial basis function (RBF) covariance kernel that defines the covariance between log firing rates at two time points *t*_1_, *t*_2_:

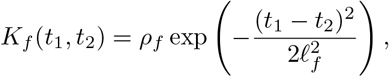

where *ρ*_*f*_ controls the variance and *ℓ*_*f*_ controls the length-scale (or smoothness) of the fluctuations in log firing rate. Similarly, we modeled the time-varying gain component of neural noise as:

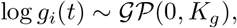

where *K*_*g*_ is an exponentiated power law (EPL) kernel that characterizes temporal correlations in the gain signal. It is defined as:

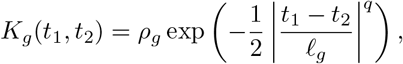

where *q* denotes the power law exponent that determines the decay of correlations at long timescales. The neural firing rate can then be expressed as the product of the stimulus-dependent drive and the stimulus-independent gain:

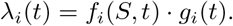

To transform firing rates into a space in which Euclidean distances between population representation vectors correspond to perceptual discriminability, we used the following transformation previously derived for the modulated Poisson process [28]:

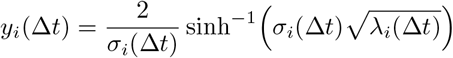

where *λ*_*i*_(Δ*t*) refers to the mean spike count of neuron *i* in time bin Δ*t*:

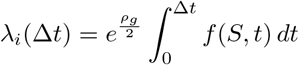

and *σ*_*i*_(Δ*t*) is the contribution of firing rate variability:

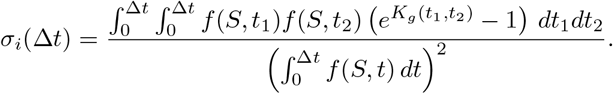

Population curvature was computed by first defining a sequence of unit displacement vectors **v**_Δ*t*_ from successive neural embeddings **y**_Δ*t*_, **y**_Δ*t*−1_ ∈ ℝ ^*N*^ :

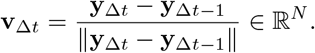

Population curvature at time window Δ*t* was then calculated as the angle between successive displacement vectors, computed from their dot product:

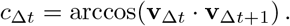

For the experiments in Fig. 7, we simulated a population of *N* = 100 units for a duration of *T* = 2.2 *s* at a resolution of *dt* = 10^−3^ *s*. To generate the stimulus-dependent tuning functions *f* (*S, t*), we set *ρ*_*f*_ = 3, which determined the range of signal amplitudes. Tuning function timescales were chosen in the range *ℓ*_*f*_ ∈ [10^−3^, 2], corresponding to the abscissa in Fig. 7d,e. To control the amplitudes of the stimulus-independent gain signals, the marginal variance was chosen to be *ρ*_*g*_ = 0.1. To study the effect of noise timescales on trajectory curvature, we chose *ℓ*_fast_ = 2 × 10^−3^ to generate fast gain signals and *ℓ*_slow_ = 2 × 10^−2^ for slow gain signals and fixed *q* = 2. Since both the length-scale *ℓ*_*g*_ and the power-law exponent *q* control the temporal smoothness of the noise, we conducted additional simulations fixing *ℓ*_*g*_ = 2 × 10^3^ while varying *q* ∈ [0.2, 2]. We found no qualitative differences in the results, indicating that the specific choice of hyperparameter used to control noise timescales is not critical.

Each dot in Fig. 7d,e represents the mean curvature 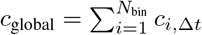 across time bins *N*_*bins*_ for a given tuning timescale *ℓ*_*f*_.

## Acknowledgements

We thank Hayoung Song and Eli Merriam for helpful comments on an early version of this manuscript. This work was supported by US National Institutes of Health grants EY032999 (R.L.T.G.), DC020088 (A.G.H.) and National Science Foundation CAREER award #2146369 (R.L.T.G.).

## Author contributions

J.X., A.G.H. and R.L.T.G. conceived and designed the study. J.X. and J.T. analyzed the fMRI data. J.X. and T.D.N. performed model simulations. J.X., A.G.H. and R.L.T.G. wrote the manuscript with input from T.D.N and J.T..

## Data and code availability

The data underlying this study are already public on OpenNeuro (https://openneuro.org/datasets/ds003020/versions/3.1.1), and the code will be made available before publication.

**Extended Data Figure 1.**
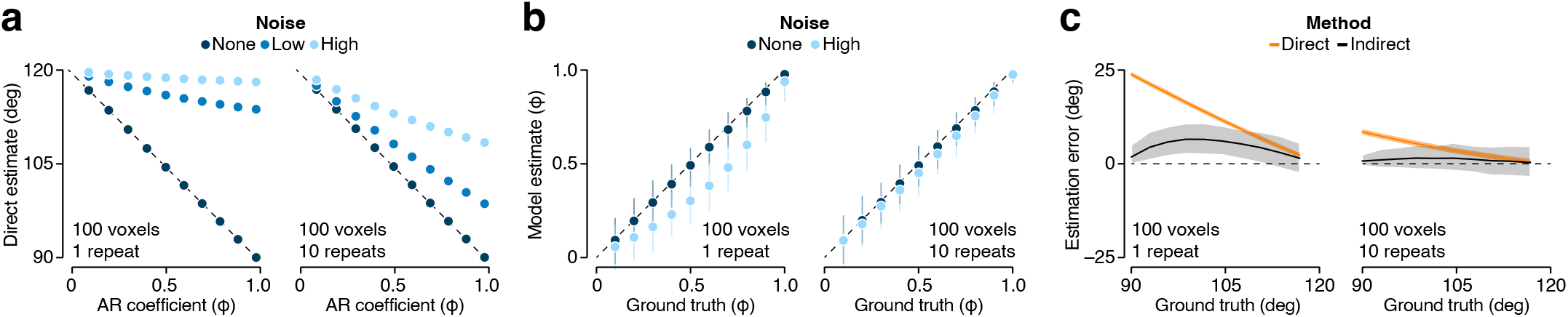
Comparing estimation errors for single-trial and trial-averaged response trajectories. (**a**) (Left) Direct curvature estimates are plotted against AR-coefficient for a set of simulated 100-dimensional trajectories, which varied in their signal (X-axis) and noise (color) properties. Symbols show the mean across 500 runs of the simulation. (Right) Same plotting conventions, now applied to trial-averaged trajectories. (**b**) Estimated representational timescale plotted against ground truth value for AR(1) response trajectories that varied in timescale (abscissa) and noise level (light vs dark colors). Symbols show the mean across 500 runs, error bars indicate the cental 95 % of the distribution. (Right) Same plotting conventions, now applied to trial-averaged trajectories. (**c**) Curvature estimation error as a function of ground truth value for two estimation methods (orange vs black), applied to a set of simulated 100-dimensional trajectories. Thick lines show the mean across 500 runs of the simulation, colored bands indicate the central 95 % of the distribution. (Right) Same plotting conventions, now applied to trial-averaged trajectories.

**Extended Data Figure 2.**
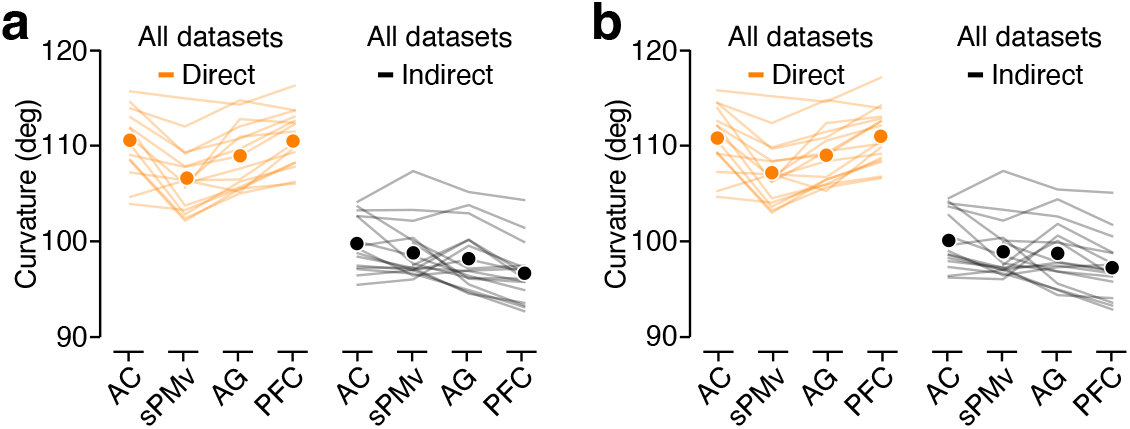
Comparison of curvature estimates across four brain regions. (**a**) Curvature estimates for all datasets (each line shows one dataset; symbols show the average). (Left) Direct method. (right) Indirect method. This is the analysis shown in the main paper. Due to differences in the size of each brain region, estimates are based on different numbers of voxels.(**b**) Same analysis, using the same number of voxels for each brain region.

**Extended Data Figure 3.**
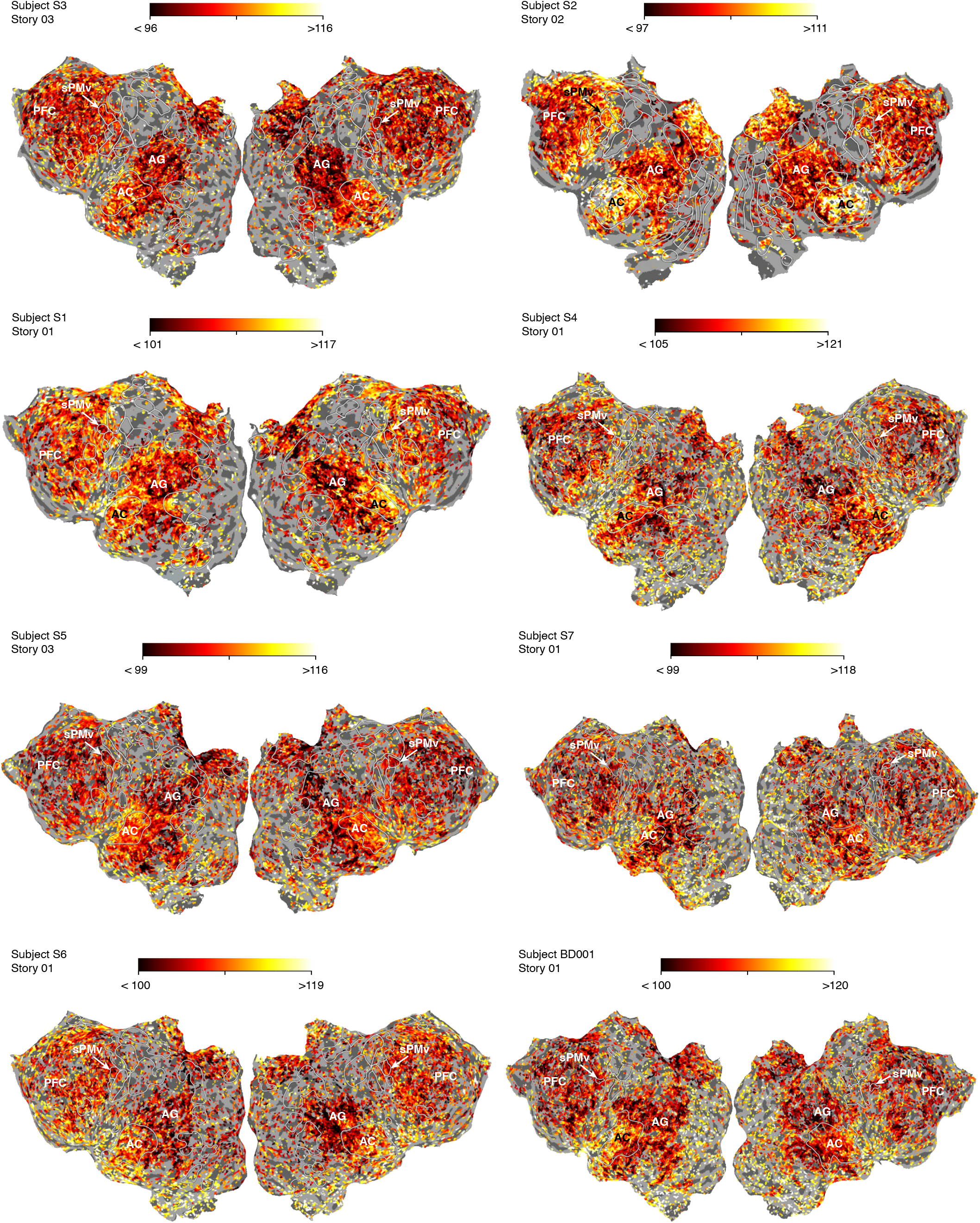
Subject-level cortical maps of implied trajectory curvature. Curvature estimates are shown separately for each subject while listening to a story. Values were computed using the same indirect AR(1)-based procedure and reliability threshold as in Fig. 3. Across subjects, implied curvature is highest in medial AC and decreases progressively both within AC and along the broader cortical hierarchy toward higher-order speech processing areas, consistent with the spatial gradient shown for the example subject in Fig. 3.

**Extended Data Figure 4.**
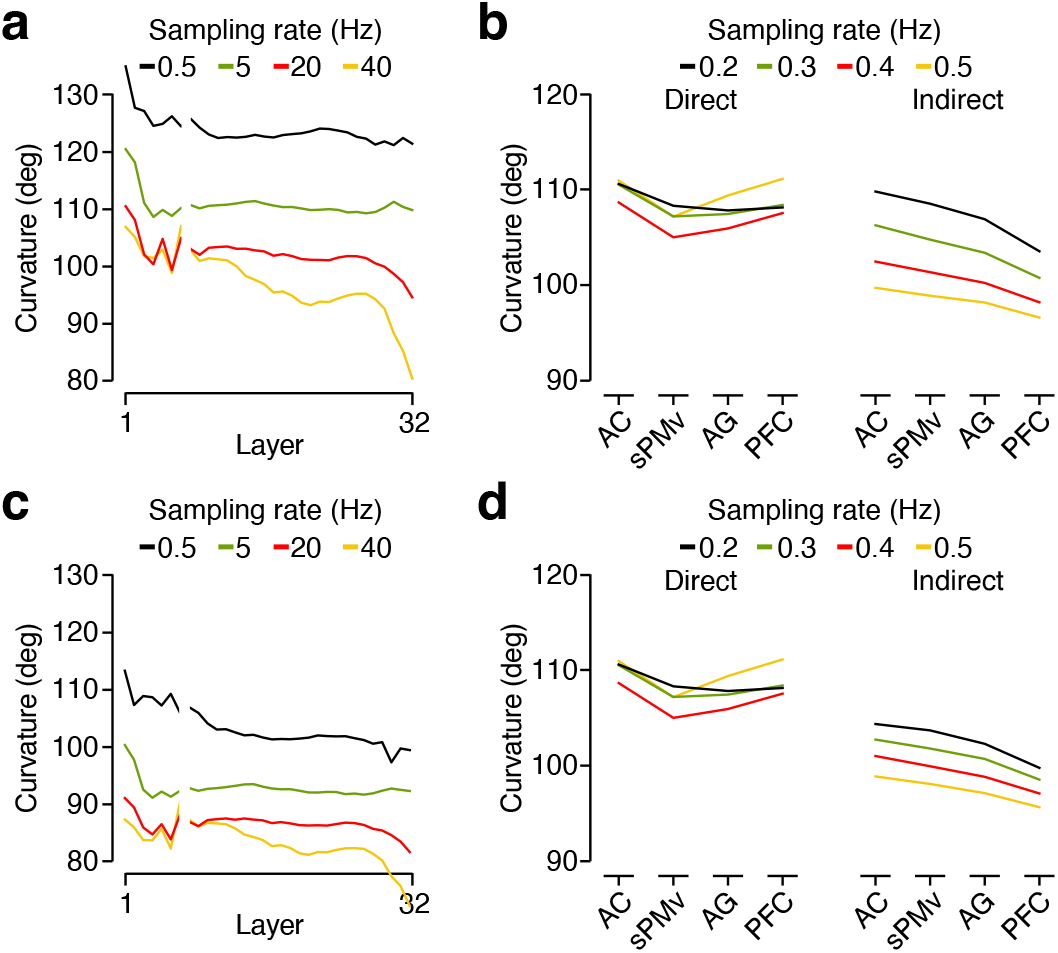
Comparison of trajectory curvature across temporal scales for different down-sampling methods. (**a**) Comparison of wavLM trajectory curvature across different sampling rates. Estimates were obtained with the direct method. (**b**) Comparison of fMRI trajectory curvature across different sampling rates. Lines show the average curvature estimates across all datasets. Left: direct method. Right: Indirect method. (**c-d**) Same as panel a-b for a down-sampling method that also includes low-pass filtering (Methods).

